# Sex-dependent effects of a gestational ketogenic diet on offspring birth and lifespan

**DOI:** 10.1101/2025.02.19.639045

**Authors:** Sarah M. Zala, Renata Santos, Eva Strasser, Alice Schadde, Sarah Kugler, Verena Strauss, Anna Kübber-Heiss, Diana Zala

**Author notes:** Corresponding authors: Diana Zala,. Université Paris Cité, Institute of Psychiatry and Neuroscience of Paris (IPNP), INSERM U1266, Dynamics of Neuronal Structure in Health and Disease, 75014 Paris, France; Sarah M. Zala,. Konrad Lorenz Institute of Ethology, University of Veterinary Medicine, Vienna, Savoyenstrasse 1a, 1160 Vienna, Austria; Renata Santos,. Université Paris Cité, Institute of Psychiatry and Neuroscience of Paris (IPNP), INSERM U1266, Dynamics of Neuronal Structure in Health and Disease, 75014 Paris, France. Equal contribution (Sarah M. Zala, Renata Santos).

## Abstract

Low-carbohydrate, high-fat ketogenic diets (KDs) are used to treat drug-resistant epilepsy, and other potential benefits for treating neurological disorders, metabolic syndrome, and cancer are being explored. In addition to these and other medical applications, KDs have also become popular for rapid weight-loss and enhancing athletic performance. However, the effects of exposing developing offspring to KDs during pregnancy (gestational KD) are poorly understood, and especially their long-term health consequences. In this study, we investigated the effects of a partial gestational KD during the second half of pregnancy in mice and assessed the consequences on the offspring over their entire lifespan compared to offspring exposed to a control diet. We found that a gestational KD significantly reduced dams’ litter size and litter mass and altered the sex ratio at birth, reducing the proportion of female offspring, which also had lower body mass early in their life. In contrast, male offspring exposed to a gestational KD suffered a significantly reduced lifespan and a late-onset increase in body mass. We found no evidence that our KD diet influenced adult offspring behavior (locomotion, anxiety, depression, circadian rhythms, food and water consumption) or reproduction. These findings highlight the potential of even a partial maternal exposure to a KD to have surprisingly detrimental effects on offspring health and longevity, and thus raising concerns about its use during pregnancy.

## Introduction

Ketogenic diets (KD) have been used for over a century to treat drug-resistant epilepsy and they are currently being investigated for a variety of potential therapeutic benefits, including treating neurodegenerative diseases, psychiatric disorders, metabolic syndrome, and cancer (1–4). Recently, KDs have gained popularity for weight loss and enhancing athletic performance, despite the limited understanding of their physiological and long-term health consequences in humans (5).

KDs, characterized by low carbohydrate, moderate protein, and high fat intake, induce a metabolic switch known as ketosis, in which ketone bodies replace glucose as the primary energy source (6,7). This metabolic shift mimics a physiological state that normally occurs during fasting, resulting in elevated serum ketones while maintaining moderate to normal blood glucose and physiological pH levels (8). Ketone bodies — acetoacetate, β-hydroxybutyric acid (BHB), and acetone — are products of fatty acid degradation in the liver. Acetoacetate is the primary ketone produced, which is subsequently converted into BHB and volatile acetone before being released into the bloodstream. Increased BHB concentration in the blood is thus used as a marker of ketosis (9). In target tissues, acetoacetate and BHB are efficiently metabolized into acetyl-CoA within mitochondria, fueling the tricarboxylic acid cycle and ATP synthesis (7). However, this metabolic shift, with its drastic reduction in glycolysis, may affect tissues that rely on glycolytic ATP for specific functions, such as neurons in the brain (9–12). Besides its metabolic role, BHB acts as a direct and indirect signaling molecule, for instance in epigenetics and catabolism regulation (13). Overall, a KD induces a metabolic reprogramming that is accompanied by pleiotropic physiological effects on the organism. The problem is that prolonged glucose restriction can potentially cause lasting and potentially irreversible alterations, particularly during *in utero* development.

The impact of a gestational KD on offspring development, health, and survival remains largely unexplored and so the results can be briefly summarized. One study reported that sons of epileptic mothers treated with a moderate KD during pregnancy displayed normal development at birth (5,14). In contrast, studies on gestational diabetes mellitus revealed an inverse correlation between maternal gestational ketone levels and the intelligence quotient of offspring (9,15). Studies in rodents showed that a gestational KD during the entire gestation period affects offspring as early as embryonic pre-implantation stages (16–19) and induces postnatal developmental delays, structural brain differences, and behavioral abnormalities that persist into adulthood (20–22). Conversely, other research found that gestational KD exposure reduced depression- and anxiety-like behaviors and increased sociability, without altering oxytocin expression (23,24). Lactation is also a critical period, as adverse effects have been observed in both lactating rats and their offspring (20). Additionally, a severe case of starvation ketoacidosis in a breastfeeding mother has been documented (25). These findings in humans and rodents underscore the complexity of gestational KDs, emphasizing the need for further research in this area.

In the present study, we investigated the effects of a gestational KD limited to the organogenesis period (late gestation) and assessed its impact across the entire lifespan of the offspring. Our findings reveal that even a partial, 10-day maternal metabolic shift to ketosis can have sex-specific detrimental effects during critical life stages. Female offspring were adversely affected at birth, while male offspring had a reduced longevity and an increased body mass late in life.

## Results

### Induction of a partial gestational ketosis with a 10-day ketogenic diet

Previous studies showed that a prolonged KD exposure before, during, and after gestation reduced fertility, delayed embryonic development, and induced organ-specific changes in the progeny (19,26). Additionally, *in vitro* studies reported that physiological concentrations of ketones induced epigenetic modifications in blastocysts, impaired implantation, and caused female-specific developmental perturbations (17,18). To investigate the effects of a gestational ketosis specifically at late embryogenesis, a KD administration was initiated at gestational day (G)8.5, when organogenesis begins, after the fetal neural plate and heart tube had already formed. The diet was terminated at G18.5 to prevent potential adverse effects during lactation. The KD used in this study comprised 5% carbohydrates, 11% protein, and 84% fat. This carbohydrate proportion closely mirrors human KD formulations but is significantly higher than the ultra-low carbohydrate content (<1%) commonly used in murine studies. This partial and mild dietary intervention allowed us to focus on the effects of gestational ketosis on organogenesis while minimizing the broader disruptions associated with prolonged or more extreme dietary regimens.

Blood glucose and BHB levels were monitored throughout the gestation, with measurements taken pre-diet (G7.5 and G8.5), during the diet (G10.5, G12.5, and G18.5), and post-diet (G19.5, after reintroduction of the control diet). As expected, during the KD period the KD group exhibited a moderate reduction in blood glucose levels (ANOVA with repeated measurements: n=17, time: F=18.50, p<0.00001; time*diet: F=2.07, p=0.008). This reduction was statistically significant at G10.5 (T-test: n=8, T=0.79, p=0.016; Fig 1A). In contrast, BHB levels significantly increased during the KD period, confirming a state of ketosis (ANOVA with repeated measurements: time: n=16, F=12.396, p<0.0001; time*diet: n=16, F=10.868, p<0.0001; post-hoc test: p<0.001 at G10.5, G12.5, and G18.5; Fig 1B). These results confirm that a gestational ketosis was successfully induced. As expected, the body mass of pregnant females in both groups increased throughout gestation and decreased after delivery. However, at G18.5 the KD mice showed a significantly lower body mass compared to the CD group (ANOVA with repeated measurements: n=16, time: F=398.24, p<0.00001; time*diet: F=77.40, p<0.00001; post-hoc test: p=0.003 at G18.5; Fig 1C). This difference resolved post-delivery.

**Fig 1.**
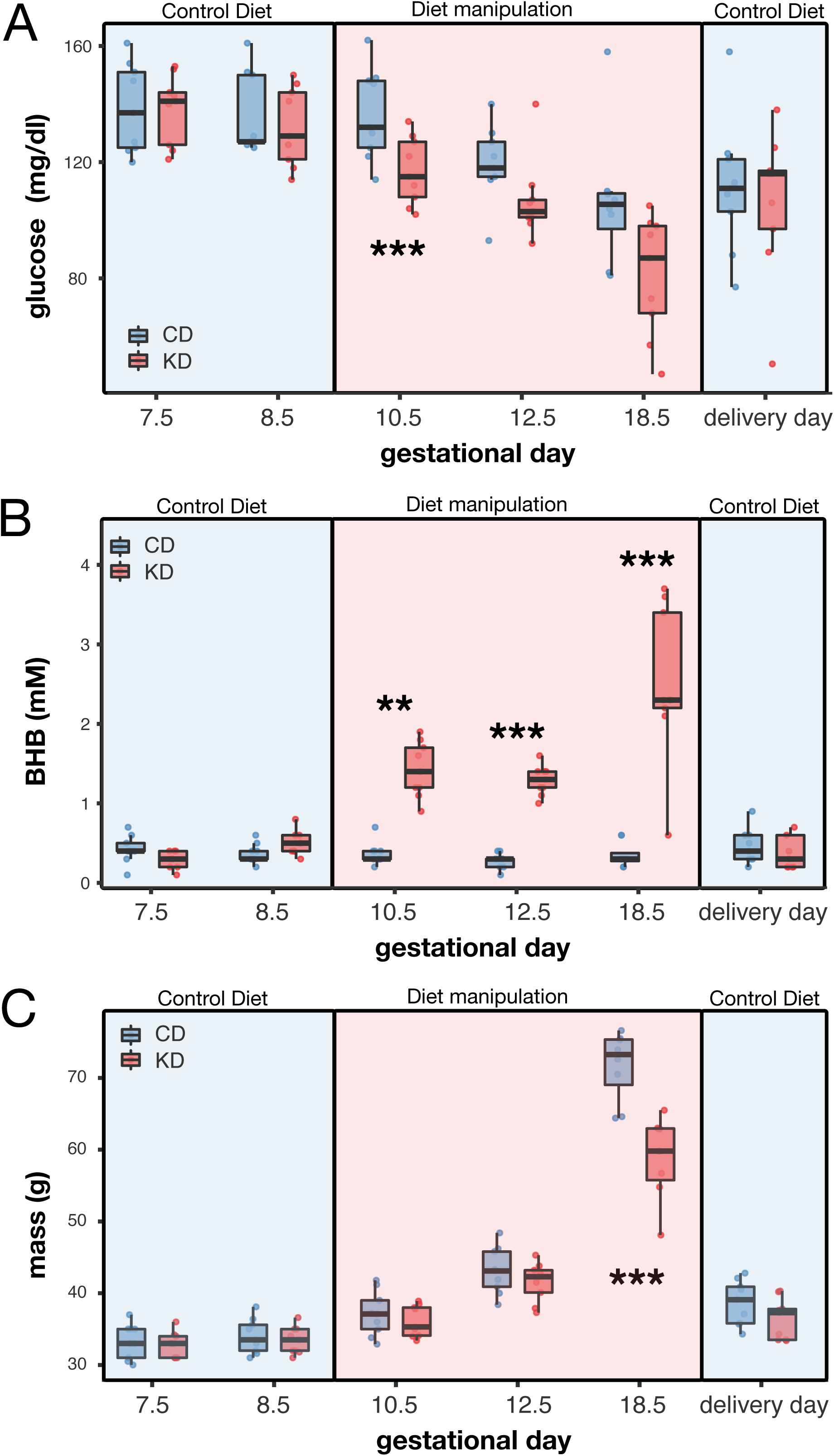
Effects of a partial KD in dams’ glucose and ketone levels and body mass during gestation. (A-C) Box plots showing the dynamics of (A) blood glucose, (B) BHB, and (C) body mass in dams fed either a control diet or ketogenic diet (**p<0.01, ***p<0.001). Box plots include all values with outliers, median, individual values, first and last quartiles, whiskers drawn within the 1.5 interquartile range value.

In summary, our partial KD effectively induced ketosis, maintained physiological glucose levels, although reduced compared to a control diet, and resulted in a transient reduction in maternal body mass during late gestation.

### A partial gestational KD influenced litter size, offspring sex ratio, and pup body mass

Our gestational KD had a striking effect on the litter size and sex ratio of the offspring at birth. Litter size was significantly reduced in the KD group compared to the CD offspring (independent samples T-test: n=9 for KD and n=10 for CD, T=3.29, p=0.005; Fig 2A). Notably, one female in the KD group failed to deliver, and this animal was conservatively excluded from the analysis. Unexpectedly, the number of female pups per litter was significantly reduced in the KD group, while the number of male pups remained unchanged (independent samples T-test: n=9 for KD and n=10 for CD; females: T=3.04, p=0.007; males: T=0.15, p=0.885; Fig 2B). The offspring sex ratio in the CD group adhered to a Mendelian distribution (74 females and 66 males; 52.9% females; Fig 2C). However, in the KD group, females comprised only 39.6% of the offspring (38 females and 58 males; χ² test: p=0.045). These findings suggest that female embryos were more vulnerable to gestational ketosis, potentially leading to increased embryonic lethality.

**Fig 2.**
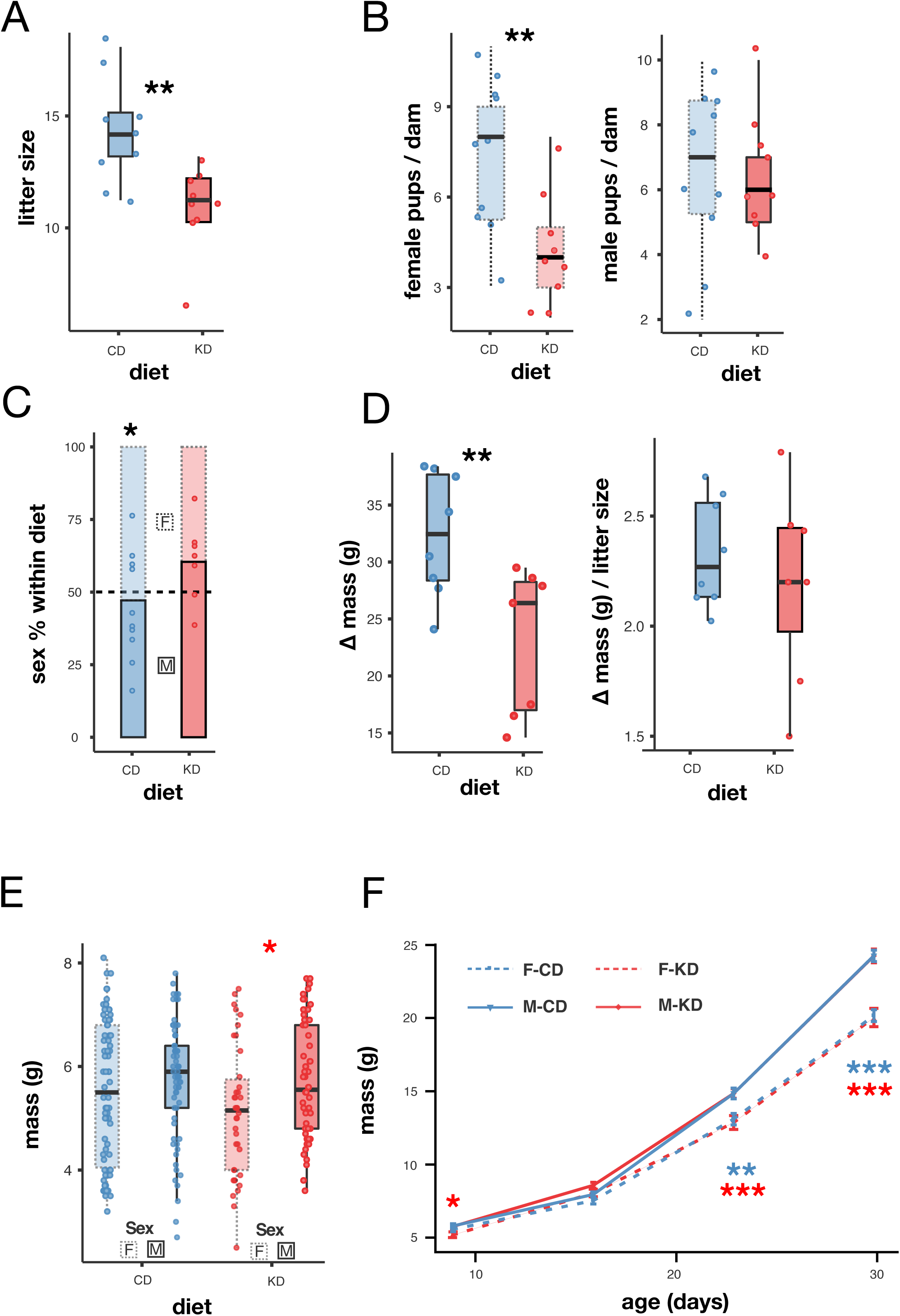
A partial gestational KD impacted female, but not male, pups. (A) Litter size by gestational diet. (B) Number of female (left) and male (right) pups per litter by diet. (C) Offspring sex ratio by diet. (D) Maternal body mass difference between G18.5 and post-delivery (G18.5-G19.5) by diet (left) and estimated mean pup mass calculated by dividing the maternal mass difference by the litter size (right). (E) Pup body mass at P9 by diet and sex. (F) Dynamics of pup body mass during the first month by diet and sex (mean±SEM). Box plots include all values with outliers, median, individual values, first and last quartiles, whiskers drawn within the 1.5 interquartile range value. Statistics A, B, D, and E: Student’s T-test; C: χ² test, F: repeated measures ANOVA. Significance levels: *p<0.05, **p<0.01, ***p<0.001.

Maternal body mass pre-delivery (G18.5) and post-delivery (G19.5) was compared to estimate the total delivery mass as a proxy for litter mass. A significant reduction in this mass was observed in the KD group compared to the CD group (T-test: n=7 for KD and n=8 for CD, T=3.06, p=0.009; Fig 2D). When normalized by litter size, the average mass per pup showed a smaller and non-significant reduction in the KD group (independent samples T-test: n=7 for KD and n=8 for CD, T=0.78, p=0.450; Fig 2D). These results contain mean values for both males and females. To further evaluate offspring growth, pup body mass was monitored weekly from postnatal day nine (P9) to weaning. While male and female pups in the CD group had comparable body masses at P9 (independent samples T-test: T=1.033, p=0.302; n=65 for males and n=74 for females; Fig 2E), female pups in the KD group were significantly lighter than males at this stage (independent samples T-test: T=2.32, p=0.023; n=58 for males and n=38 for females; Fig 2E). This finding suggests that female pups from KD dams were already smaller at birth and that this difference persisted for at least nine days, despite the fact that during postnatal lactation the mothers were no longer in ketosis. At P16 female body mass in the KD group did not differ from male body mass (T=1.81, p=0.074; Fig 2E). However, starting at P23 a significant sexual dimorphism in body mass was evident in both dietary groups, with males consistently weighing more than females, as expected (ANOVA with repeated measures: age p>0.0001, age*sex p<0.0001, age*diet p=0,012, Post Hoc Test with Holm correction at P23 and P31, sex: p <0.0001, Fig 2F).

In summary, a partial gestational ketosis had detrimental effects on female offspring by reducing their number at birth and their initial body mass, whereas male offspring appeared unaffected.

### A partial gestational KD had no lasting impact on offspring metabolic profile

To avoid potential confounding effects of extreme litter sizes, two outliers from the CD group (litter sizes of 16 and 17 pups) were excluded from further analysis. Glucose and BHB levels were measured in adult offspring at four months of age. All offspring displayed glucose levels within the normal glycemic range and low BHB levels, as expected, indicating no lasting metabolic alterations from gestational KD exposure (ANOVA: sex*diet, for glucose p=0.552, F=0.353; for BHB p=0.668; F=0.186; n=18 and n=17; Fig 3A). In addition, there was no difference in body mass (ANOVA between subjects effects: sex*diet: p=0.734, F=0.115, df=228; Fig 3B). Finally, no differences were observed in food and water consumption (ANOVA: sex*diet, food: p=0383, F=0.383; water: p=0.965, F=0.002 n=12; Fig 3C). These findings suggest that a partial gestational KD exposure did not result in lasting metabolic effects on adult offspring.

**Fig 3.**
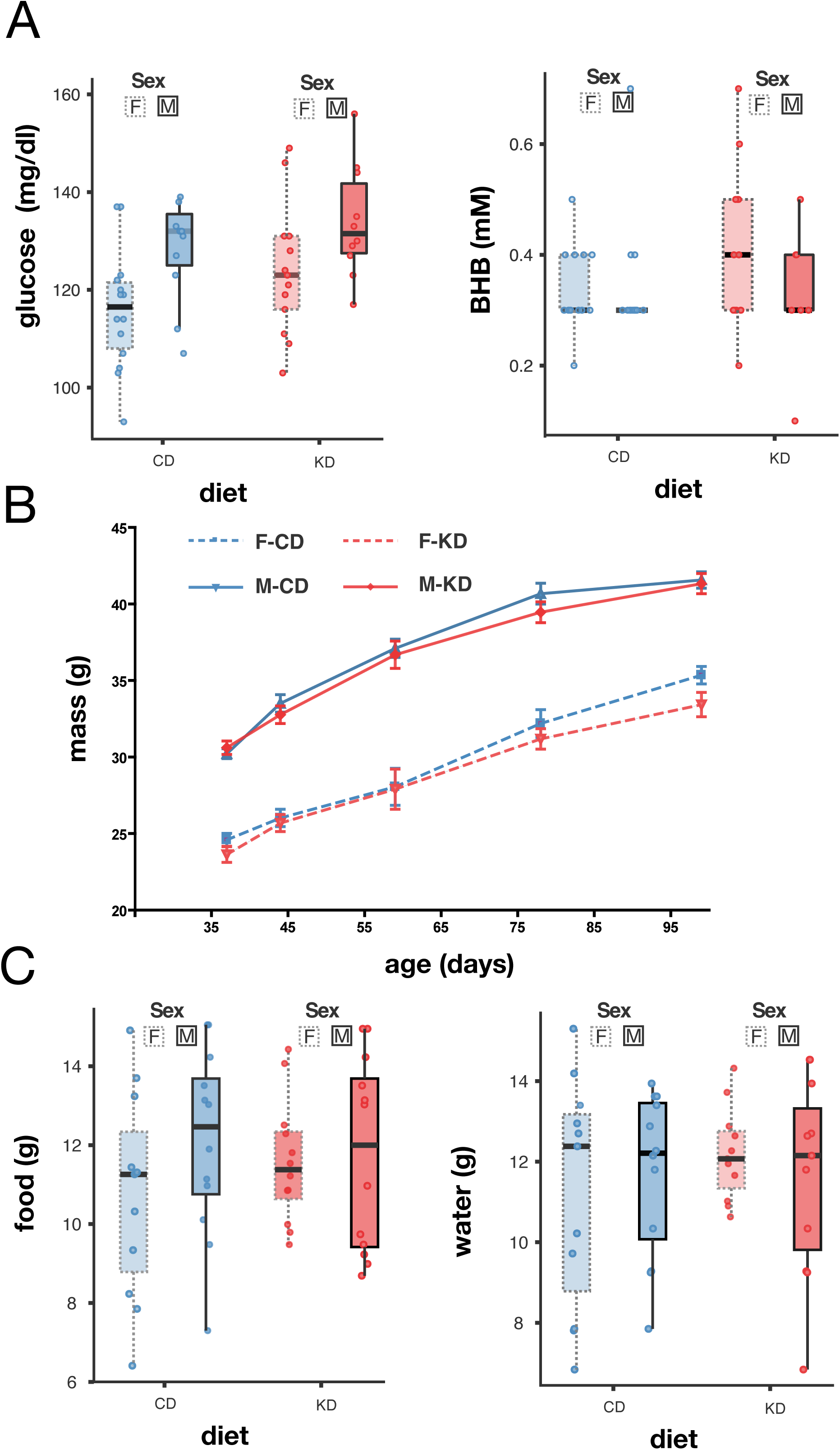
A partial gestational KD did not affect metabolism, body mass, nor food and water intake. (A) Blood glucose (right) and blood BHB levels (left) in adult offspring by diet and sex. (B) Body mass dynamics across time by diet and sex. (C) Total food (left) and water consumed over 78 h. Box plots include all values with outliers, median, individual values, first and last quartiles, whiskers drawn within the 1.5 interquartile range value. Statistics A-C: ANOVA, p>0.05.

### A partial gestational KD had no lasting impact on offspring anxiety- and depressive-like behavior

In a previous study, Sussmann *et al*. (23) reported that a full gestational KD in CD-1 mice reduced susceptibility to anxiety and depression, as assessed using the open field test (OFT), elevated plus maze (EPM), and forced swim test (FST), compared to controls on a standard diet. In the present study, we evaluated spontaneous activity and found no differences in the total distance traveled in the OFT (ANOVA: sex*diet p=0.337, F=0.944, n=12) or the time spent in the center of the arena (ANOVA: sex*diet p=0.205, F=1.658, n=12; Fig 4A). Similarly, the EPM test revealed no significant differences in the time spent in the open or closed arms (ANOVA: sex*diet p=0.872, F=0.026, n=12; Fig 4B). Additionally, depression-like behavior assessed via the FST also showed no significant differences between sex or diet groups (ANOVA: sex*diet p=0.178, F=1.863, n=12; Fig 4C).

**Fig 4.**
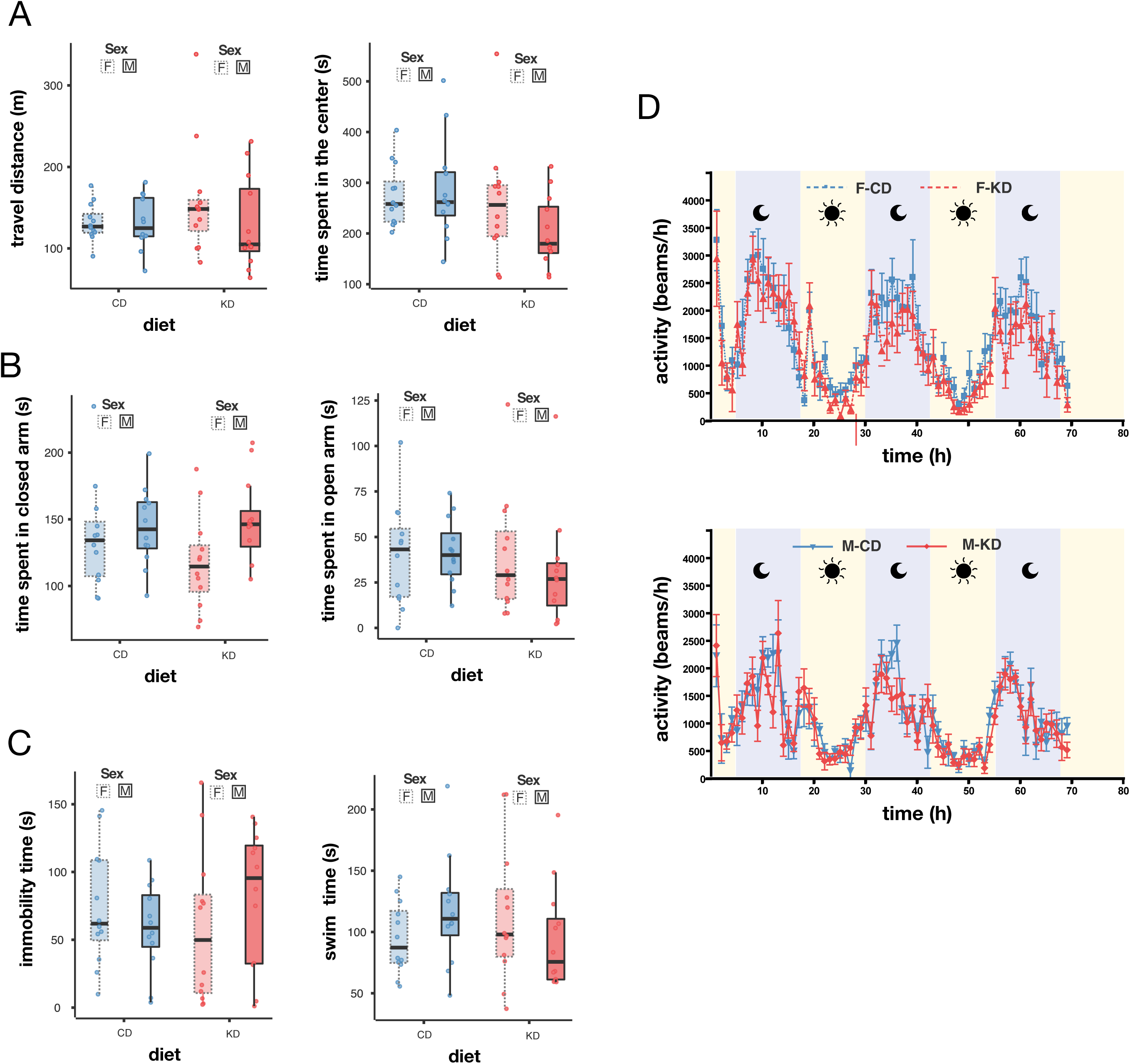
A partial gestational KD did not affect adult offspring spontaneous locomotion, susceptibility to anxiety and depression, nor circadian rhythms. (A) Total distance traveled during 1 hour in the open field test (left) and time spent in the center (right) by diet and sex. (B) Time spent in the closed arm (left) and the open arm (right) during a 5-minute elevated plus maze assay. (C) Time spent immobile (left) or actively struggling to swim (right) during a 5-minute forced swim test. (D) Locomotor activity in the light and dark cycles. Box plots include all values with outliers, median, individual values, first and last quartiles, whiskers drawn within the 1.5 interquartile range value. Line graphs (D) shows mean±SEM. Statistics A-C: Anova p>0.05.

### A partial gestational KD had no lasting impact on circadian rhythms

Dietary interventions, such as high-fat diets and caloric restriction, can disrupt the oscillatory expression of genes, thereby altering circadian rhythms (27,28). E.g., KDs increase ketone body production, mimicking a fasting state and causing shifts in circadian phases. The establishment of circadian rhythms begins during late embryonic development and is influenced by maternal factors (29–31). To investigate whether maternal ketosis induced circadian changes in our model, we evaluated night-day rhythms. Our results showed no significant differences between gestational KD and control animals (Fig 4D).

In conclusion, a 10-day gestational ketogenic diet caused no significant behavioral abnormalities related to anxiety, depression, spontaneous activity or circadian rhythms in the adult offspring.

### A partial gestational KD had no lasting impact on offspring reproductive success

To measure offspring fertility and reproductive success, we bred 22 gestational CD and 10 gestational KD pairs. All 32 females successfully gave birth; however, two gestational KD females and one CD female cannibalized their offspring post-delivery. The latency to give birth differed significantly between the groups, with the gestational KD group delivering at 21.5±1 days and the CD group at 25.5±7 days after pairing (T-test with unequal variances: T=-2.6, df=24.3, p=0.016; Fig 5A). Despite this difference in latency to give birth, no significant differences in reproductive success were observed when offspring were weaned (mean number of weaned offspring: KD=10.2±6, CD=11.3±5; T-test: T=0.5, df=30, p=0.59). Additionally, the offspring sex ratio followed a Mendelian distribution in both groups (CD: 115 females and 133 males; KD: 46 females and 56 males; χ² test: p=0.828; Fig 5B).

**Fig 5.**
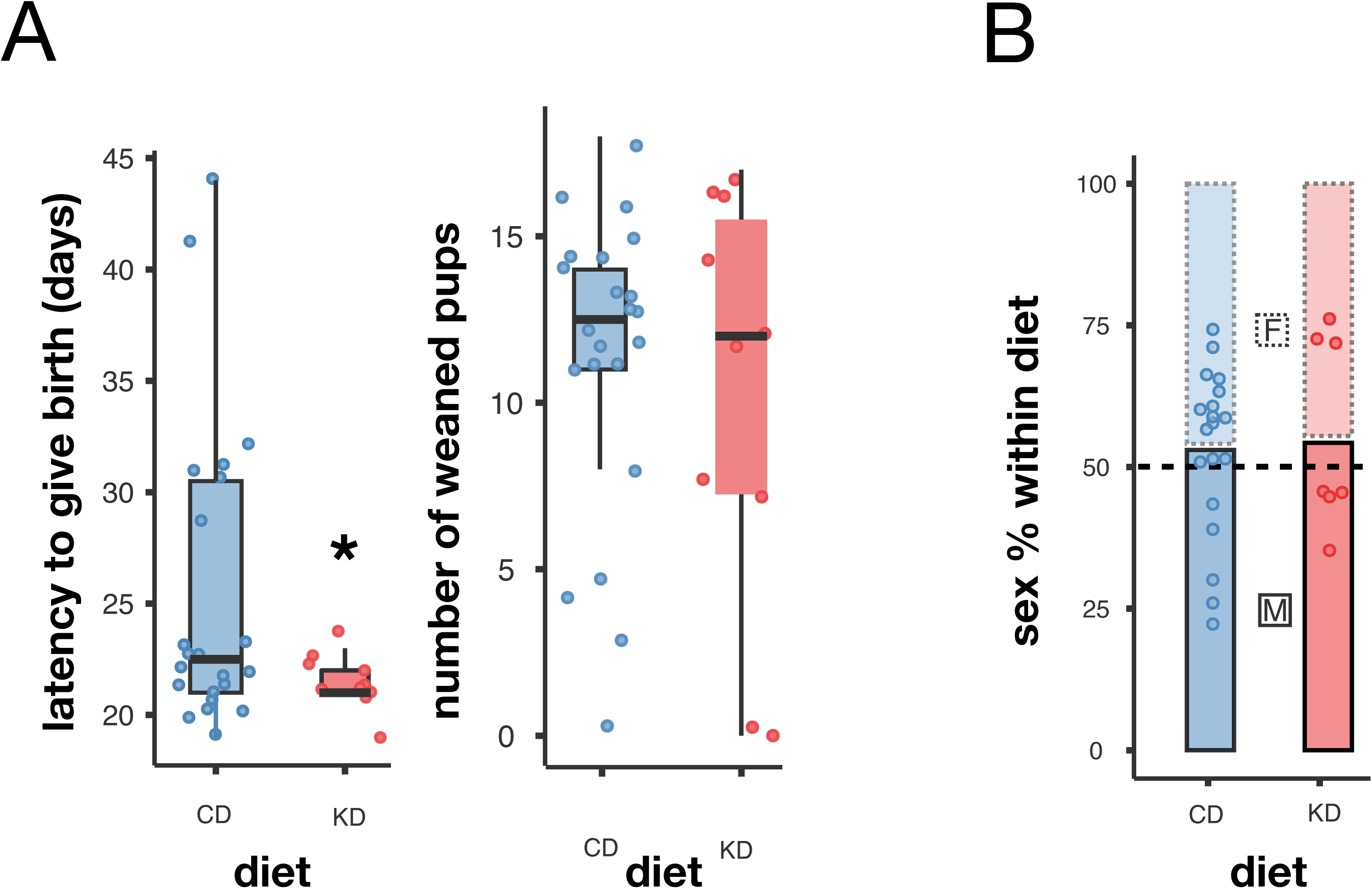
A partial gestational ketogenic diet had no lasting impact on offspring’s reproductive success. (A) Latency to give birth (left) and number of weaned pups (right) by diet. (B) Offspring sex ratio by diet. A: T-test, p<0.05; B: χ² test test, p>0.05

### A partial gestational ketogenic diet resulted in late-life increased body mass and shortened lifespan of male offspring

We found that a maternal KD during gestation resulted in reduced female offspring body mass shortly after birth (Fig 2E), though this difference resolved by P16 (Fig 2F). No further differences in female body mass were detected throughout their life until their natural death (Fig 3B and data not shown). No differences in body mass were observed in KD male offspring until approximately 2.2 years of age (Figs 2E-F, 3B, 6A) when a significant increase in body mass was found, which persisted until their natural death (Fig 6A).

**Fig 6.**
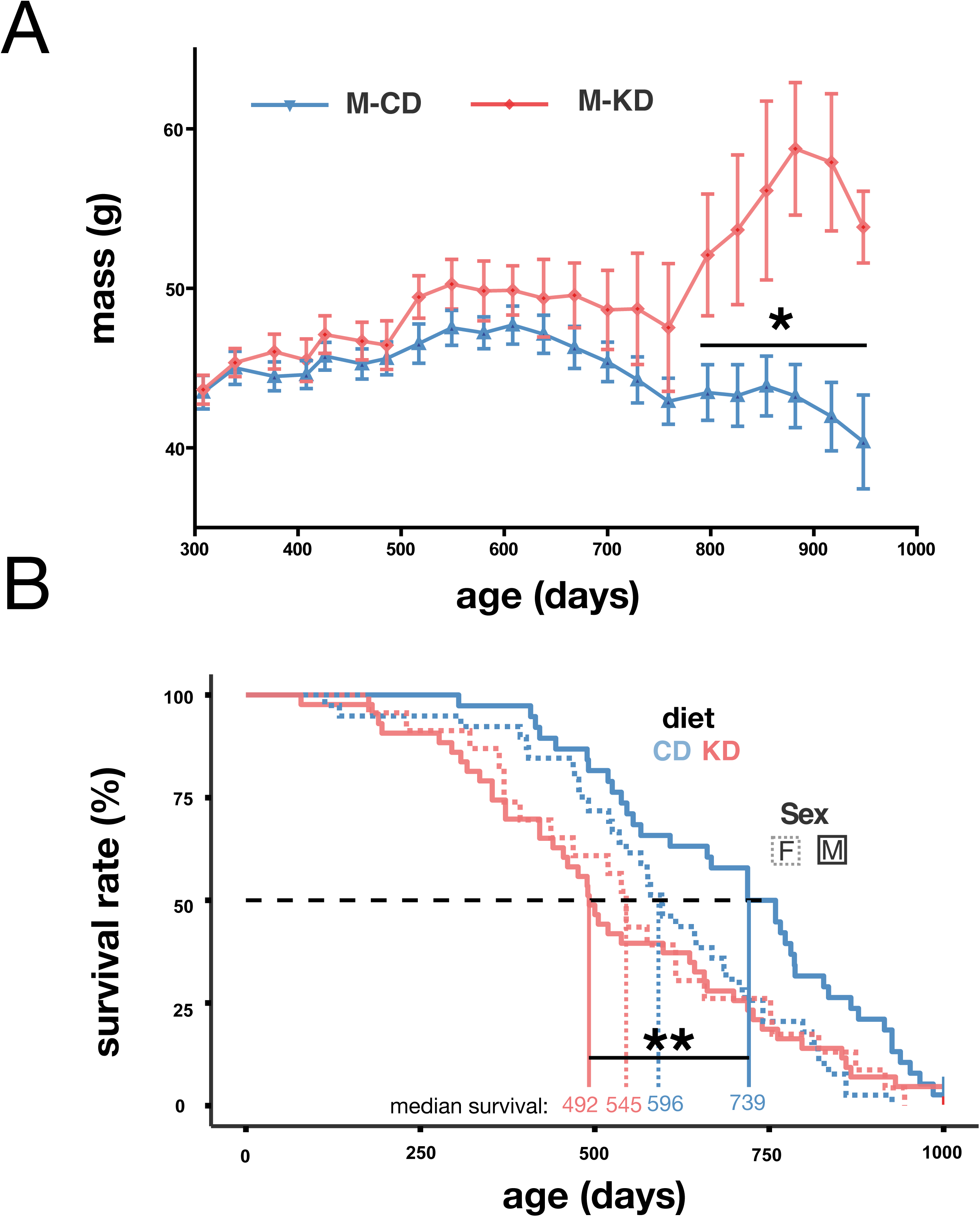
A partial gestational KD impacted male lifespan and mass during late-life. (A) Body mass dynamics across time by diet: Repeated ANOVA, p<0.05. (B) Survival rate of the offspring by sex and diet: Log-rank test: χ²=7.47, p=0.006.

We assessed offspring longevity in two cohorts (one in France and one in Austria) with two different husbandry systems, one specific pathogen free (SPF) and one conventional (see Materials and Methods). The long-term survival of offspring did not significantly differ between the French and the Austrian cohorts (Log-rank test: χ²=2.14, p=0.144), which allowed pooling these data (Fig 5B). No sex-bias was detected in the overall longevity (Log-rank test: χ²=1.285, p=0.257). However, median survival was significantly reduced in the KD offspring (512 days) compared to controls (661 days) (Log-rank test: χ²=7.47, p=0.006). Sex- and diet-specific analyses revealed that male KD offspring had a significantly reduced median survival of 247 days compared to controls (KD: 492 days; CD: 739 days; Log-rank test: χ²=9.459, p<0.0021, Fig 6B). In contrast, female survival was only slightly affected by the diet, with median survival differing by 51 days (KD: 545 days; CD: 596 days; Log-rank test: χ²=0.38602, p=0.2344).

In summary, a partial gestational ketogenic diet had lasting effects on male offspring, including a late-onset increase in body mass and a reduction in lifespan, but these effects were not found in females.

### A partial gestational ketogenic diet did not alter pathological outcomes in offspring

We performed pathological examinations on 66 mice (CD: n=40, KD: n=26), including 21 that were euthanized due to health issues and others found dead in their cages. The examined cohort included 21 males and five females from the KD group, and 19 males and 21 females from the CD group (Table 1). Parasitological analysis identified oxyurid infections in four mice (KD: n=1, CD: n=3).

**Table 1.** Offspring pathology: Type of neoplasia and level of malignancy according to diet and sex. Note that some individuals had more than one type of neoplasia. Each row shows the number (and % total) of mice with the specific disease.

|  |  | Control Diet |  |  | Ketogenic Diet |  |  |
| --- | --- | --- | --- | --- | --- | --- | --- |
| Type of neoplasia | Level of malignancy | Females<br>(n=21) | Males<br>(n=19) | Total<br>(n=40) | Females<br>(n=5) | Males<br>(n=21) | Total<br>(n=26) |
| Lymphoma | Malignant | 10 | 6 | 40% | 2 | 11 | 50% |
| Pancreatic islet cell carcinoma | Malignant | 2 | 1 | 7.5% | 2 | 1 | 11.5% |
| Pulmonary carcinoma | Malignant | 5 | 3 | 20% | 3 | 0 | 11.5% |
| Hemangiosarcoma | Malignant | 1 | 4 | 12.5% | 0 | 1 | 4% |
| Uterine carcinoma | Malignant | 1 | 0 | 2.5% | 1 | 0 | 4% |
| Squamous cell carcinoma | Malignant | 3 | 1 | 10% | 0 | 0 | 0% |
| Fibrosarcoma | Malignant | 0 | 2 | 5% | 0 | 0 | 0% |
| Renal carcinoma | Malignant | 0 | 2 | 5% | 0 | 0 | 0% |
| Mammary carcinoma | Malignant | 2 | 0 | 5% | 0 | 0 | 0% |
| Hepatic adenoma | Benign | 1 | 0 | 2.5% | 0 | 4 | 15% |
| Pulmonary adenoma | Benign | 0 | 1 | 2.5% | 0 | 1 | 4% |
| Renal adenoma | Benign | 0 | 1 | 2.5% | 0 | 1 | 4% |
| Adenoma of Harderian gland | Benign | 0 | 1 | 2.5% | 0 | 1 | 4% |
| Trichoepithelioma | Benign | 0 | 1 | 2.5% | 0 | 0 | 0% |
| Sebaceous gland adenoma | Benign | 1 | 0 | 2.5% | 0 | 0 | 0% |
| Non-neoplastic | Non-neoplastic | 1 | 3 | 10% | 1 | 7 | 31% |

One or more neoplastic lesions were observed in 73% (n=19/26) of KD offspring, with 69% (n=18) presenting malignant neoplasms. Among these, multicentric lymphoma was the most common (50%), followed by pancreatic islet cell carcinoma (11.5%), pulmonary carcinoma (11.5%), hemangiosarcoma (4%), and uterine carcinoma (4%). Four KD offspring also exhibited hepatic adenomas, while isolated cases of pulmonary adenoma, renal adenoma, and Harderian gland adenoma were noted. Seven KD mice died due to non-neoplastic causes, such as pneumonia or enteritis.

One or more neoplastic lesions were also very frequent in the CD offspring, affecting 90% (n=36/40), with 85% (n=34) developing malignant neoplasms. Multicentric lymphoma was again predominant (30%, n=12), followed by lymphomas in the spleen or lymph nodes (10%), pancreatic islet cell carcinoma (7.5%), pulmonary carcinoma (20%), hemangiosarcoma (12.5%), squamous cell carcinoma (10%), and fewer cases of fibrosarcoma, renal carcinoma, uterine carcinoma, and mammary carcinoma. Benign neoplasms were identified in two CD mice, including pulmonary adenoma, hemangioma, and sebaceous gland adenoma. Non-neoplastic causes of death were recorded in four CD offspring.

No significant differences were found between dietary groups or sexes in the prevalence of lymphoma, lung carcinoma, or pancreatic carcinoma (Table 2). Other neoplastic lesions were rare and not statistically compared.

**Table 2.** Logistic regression table: Effects of diet and sex on diagnosed neoplastic lesions. There are no significant differences between the dietary groups and sex in the occurrence of lymphoma, lung carcinoma or pancreatic carcinoma.

| Generalized Linear Model |  |  |  |  |
| --- | --- | --- | --- | --- |
| Lymphoma: |  |  |  |  |
| Coefficients: |  |  |  |  |
|  | *Estimate* | *Std. Error* | *z value* | *Pr(>\|z\|)* |
| (Intercept) | -0.7732 | 0.4935 | -1.567 | 0.117 |
| Sex | 0.6779 | 0.6592 | 1.028 | 0.304 |
| Diet | 0.8685 | 0.6592 | 1.318 | 0.188 |
| Sex:Diet | -1.1787 | 1.2078 | -0.976 | 0.329 |
| Pancreatic islet cell carcinoma: |  |  |  |  |
| Coefficients: |  |  |  |  |

|  | *Estimate* | *Std. Error* | *z value* | *Pr(>\|z\|)* |
| --- | --- | --- | --- | --- |
| (Intercept) | -2.8904 | 1.0274 | -2.813 | 0.0049 |
| Sex | 0.6391 | 1.2681 | 0.504 | 0.6143 |
| Diet | -0.1054 | 1.4510 | -0.073 | 0.9421 |
| Sex:Diet | 1.9512 | 1.8685 | 1.044 | 0.2964 |
| Pulmonary carcinoma: |  |  |  |  |
| Coefficients: |  |  |  |  |
|  | *Estimate* | *Std. Error* | *z value* | *Pr(>\|z\|)* |
| (Intercept) | -1.6740 | 0.6292 | -2.661 | 0.0078 |
| Sex | 0.5108 | 0.8114 | 0.630 | 0.5290 |
| Diet | -0.1178 | 0.8858 | -0.133 | 0.8942 |
| Sex:Diet | -16.2851 | 1769.2579 | -0.009 | 0.9927 |

## Discussion

In this study, we found that maternal exposure to a KD beginning at the organogenesis stage resulted in long-lasting, sex-specific effects on offspring health and survival. Female offspring were adversely affected at birth, whereas male offspring showed reduced survival later in life. Specifically, gestational ketosis resulted in smaller litter sizes, in reduced female pup mass, and in sex-dependent female fetal loss. Interestingly, while our gestational KD did not induce male fetal abortion, it negatively impacted aging, with KD-exposed males having a shorter lifespan.

A previous study in CD-1 mice reported that KD exposure before mating reduced fertility and litter size (26). However, in the present study, a shorter gestational KD primarily affected female offspring at birth, suggesting a complex and time-dependent impact of maternal diet on fetal development. In humans, male-biased birth ratios are well-documented, with the World Health Organization (WHO) estimating an expected ratio of 103 to 107 boys per 100 girls (https://ourworldindata.org/gender-ratio). Although the sex ratio at conception is equal, female fetuses have a higher probability of abortion, suggesting that they are slightly more vulnerable during gestation (32). Evidence from mice suggests that low-fat, high-carbohydrate diets favor female offspring (33,34), whereas high-fat diets promote a male bias (35). Female placentas are more sensitive to gene regulation influenced by dietary fat content than male placentas (36,37), potentially explaining the heightened vulnerability of female fetuses to the synergistic deleterious effects of both high fat and low carbohydrates characterized in KDs.

While KDs have been shown to benefit adult mice in reducing anxiety and depression (3,38) and a full gestational KD has been reported to improve outcomes in adult offspring (23,24), we were unable to replicate these beneficial effects. Differences in the KD composition (5% carbohydrates in our study compared to <1% in others) or the timing of diet initiation (our dietary manipulation began at late gestation) could explain these discrepancies. Crucially, our KD was administered prior to critical stages of brain development, and pregnant females exhibited a controlled ketosis, as shown by increased ketone levels while maintaining a normal glucose homeostasis.

Several studies have shown that early development can have permanent effects in mammals (39,40) and that early development can also be a key factor influencing an individual’s lifetime reproductive success (39,41). Although reproductive success was not affected in the offspring of our study, females of the gestational KD group had a shorter latency to give birth after male introduction compared to controls. This difference, however, can be explained by the high variance in the control groups (driven by a few outliers) and the smaller sample size of the KD group. Further studies focusing on behavioral phenotypes, particularly social interactions and courtship, are needed to better characterize multiple effects of a gestational KD on social behaviour.

Additionally, we observed a significant reduction in the longevity of male offspring born to KD-exposed mothers. These findings were robust across cohorts, even when mice were housed under different conditions (SPF-free versus conventional mouse facilities), adding confidence to the data. CD-1 mice are known to develop spontaneous neoplasms with age (42), and our pathological analysis confirmed that most deceased mice had at least one neoplasm. However, the gestational KD did not alter the prevalence of neoplasms. Although KDs in adult mice have been shown to extend longevity and healthspan (43), our findings suggest that a KD during gestation causes opposing effects. Interestingly, gestational exposure to high-fat diets has been associated with positive effects on brain health and aging in both male and female offspring (44).

Together, our results highlight the complex and sex-dependent effects of a gestational KD on offspring development, survival, and health. These findings underscore the need for further research to delineate the specific mechanisms underlying these outcomes and to refine dietary recommendations during pregnancy.

## Future directions

There are some limitations in our study that should be considered. First, although we detected no differences in the adult offspring behavior of the KD mice compared to the CD mice using four standard behavioral assays, we may have overlooked changes in other behaviors, such as social interactions, which should be investigated in future studies. Second, although we found differences in longevity, the reasons for the premature mortality of the KD mice are still unclear. The pathology results detected a variety of malignancies as well as benign neoplasms among the deceased mice but found no differences in the incidence of neoplasms between treatment versus control mice, despite differences in their longevity. Pathology was not conducted on all the mice that died unfortunately, and since we missed 19 mice that died before reaching 471 days of age, the pathology results are biased towards older mice. In addition, we only have data on five female offspring of the KD group and due to the small sample, the lack of sex differences of the KD group is inconclusive. Future studies are thus needed to determine the pathological causes of early mortality. Interestingly, later in life, male offspring of the KD group showed a greater body mass than the controls. While we showed that our dietary manipulation had no lasting metabolic effects on adult KD offspring at four months of age , we can not exclude that their metabolism may still be altered at very late stages of life. Their heavier mass was not due to obesity, however, as the dissected mice were not fat. The mass of mice can vary over time due to several reasons, such as water retention, stomach and intestinal contents, but also pathological conditions, such as neoplasms. As our pathological investigations did not measure factors such as ingesta or neoplasm mass, future studies are needed to understand the cause of increased body mass.

## Material and Methods

### Animals

Mice (RjOrl:SWISS, Janvier Labs) were housed in ventilated cages in a specific-pathogen-free (SPF) facility in France, with food and water provided *ad libitum*. Housing conditions were maintained at a constant temperature (23±1°C) with a 12 h light:dark cycle. An additional cohort of mice (n=95 offspring, see below) was housed in Austria under similar, although conventional husbandry with open cages standard conditions (45).

All experiments were conducted in accordance with the Council of the European Union Directive 2010/63/EU of 22 September 2010 on the protection of animals used for scientific purposes. The study was approved by the French Ethical Committee (APAFIS#27654-202103101508446 v3) and by the Ethics and Animal Welfare Committee of the University of Veterinary Medicine, Vienna (ETK-027/02/2021) in accordance with the University’s guidelines for Good Scientific Practice.

### Dietary intervention and offspring

Upon arrival, all animals were switched to and maintained on a control diet (CD: 13% fat, 20% protein, 67% carbohydrate; C1000 from Genestil). Mating was performed overnight using 6-week-old animals, and the presence of a sperm plug the following morning was designated as gestational day 0.5 (G0.5). Pregnant females were randomly assigned to either the control group or the ketogenic group, with assignment balanced based on the initial day of gestation. For 10 days, from G8.5 to G18.5, females in the KD group received a ketogenic diet (KD: 84% fat, 11% protein, 5% carbohydrate; C1084 from Genestil), while control group females continued on the CD. After weaning, offspring were transitioned to standard diets with carbohydrate content exceeding 50% (diet SAFE A04).

Offspring were randomly divided (https://www.random.org/) into two cohorts, one in France and one in Austria, and then housed under different conditions. The mice in the French cohort (n=48; 12 animals/group, equal sex ratio) were housed in groups of 4–5 individuals per cage, grouped by sex and mixed by gestational diet. In the Austrian cohort (n=95; KD females=11, KD males=31, CD females=27, CD males=26; 36–40 days old) the mice were housed socially, grouped by sex and within the same gestational diet group, with most animals housed with siblings (with a standard diet, with carbohydrate exceeding 50%, V1534, Ssniff, Germany).

### Glucose and BHB measurements

A superficial cut was made at the end of the tail and 2 drops of blood were analysed for glucose and BHB with FreeStyle Optium Neo meter (Abbott).

### Behavioral phenotyping

Behavioral phenotyping assays were conducted on 12 mice per diet and sex between three and four months of age. *Open field test (OFT):* exploratory behavior and locomotor activity were measured in open-field arenas (clear Plexiglas 40x40x40 cm^3^). Each mouse was allowed to explore a novel environment for 1 hour. The distance travelled and the time spent in the center zone (20x20 cm^2^) were recorded and analyzed using Viewpoint software (Viewpoint version 5.31.0.120). *Elevated plus maze (EPM):* anxiety-related behavior was tested in a cross-shaped maze elevated 70 cm above the floor and illuminated at 50 lux. The maze consisted of two opposing closed arms (with black Perspex walls, 20 cm high) and two open, unprotected arms. Each mouse was placed in the central platform and observed for 5 min. The time spent in the open arms, closed arms, and central platform was quantified. *Forced swim test (FST):* depression-like behavior was evaluated by placing each mouse in a transparent cylinder (height: 25 cm; diameter: 13 cm) filled with water (20 cm depth; 25°C). Each mouse was recorded for 5 min, with immobility interpreted as an indicator of depression-like behavior. *Circadian, feeding and drinking activity* were monitored simultaneously using an automated feeding/activity station (TSE Systems). Mice were placed in individual Labmaster TSE cages for 80 consecutive hours. The ambulatory and fine movement in the X dimension (total activity) was used as a parameter for spontaneous activity. Movements were detected via highly sensitive sensors embedded in the measuring platform. Data acquisition was performed on the IPNP PhenoBrain core facility.

### Reproductive success

The Austrian offspring cohort was bred at 6-7 months of age. A total of 32 mating pairs were established, with females and males matched according to their maternal diet groups (CD group: n=22; KD group: n=10). Females were introduced into the male’s cage and remained together until pregnancy was visibly confirmed. The time to birth (days from pairing to delivery) was recorded, and the offspring were counted and sexed at weaning (P23).

### Pathology

A total of 66 deceased mice from the Austrian cohort underwent comprehensive pathological assessments, including necropsy and histopathological examination. Necropsies were conducted blind to diet group allocations. During the routine pathological examination, organ samples were collected, fixed in 7% neutral buffered formalin, and embedded in paraffin wax. Tissue sections were cut to a thickness of 2.5 µm, mounted on glass slides, and stained with hematoxylin and eosin using standard protocols. When bacterial infections were suspected, microbiological testing of specific organs was performed. Parasitological examinations were routinely conducted on animals with sufficient intestinal content.

### Statistical analyses

Jamovie (version 2.3.21), SPSS (version 29.0.1.0) and R (version 4.4.2) were used for statistical analysis. Results are considered statistically significant at α<0.05. All statistical tests are two-sided. Statistical tests and results are detailed in the figure legends and table descriptions.

## Acknowledgements

We thank our research assistants B. Wernisch and T. Klaus for their help; A. Sasse, T. Leclercq and J. Forget for animal care; and G. Le Pen of the IPNP Phenobrain core facility and V. Tolle for sharing behavioral protocols. We are grateful to D. Penn, Z. Lenkei, C. Hanus and L. Lokmane for comments on the manuscript.

